# Phospho-dependent Signaling during the General Stress Response by the Atypical Response Regulator and ClpXP Adaptor RssB

**DOI:** 10.1101/2021.01.11.426222

**Authors:** Jacob Schwartz, Jonghyeon Son, Christiane Brugger, Alexandra M. Deaconescu

**Affiliations:** Laboratories of Molecular Medicine Brown University Department of Molecular Biology, Cell Biology and Biochemistry Providence, RI 02903 USA

**Author notes:** Address correspondence to: Alexandra M. Deaconescu, B.E., Ph.D., Assistant Professor of Molecular Biology, Cell Biology and Biochemistry, Brown University, 70 Ship St. G-E4, Providence, RI 02903, USA.

**Keywords:** RssB, RpoS, general stress response, ClpXP, adaptor, response regulator, Y-T coupling, SdrG

## Abstract

In the model organism *Escherichia coli* and related species, the general stress response relies on tight regulation of the intracellular levels of the promoter specificity subunit RpoS. RpoS turnover is exclusively dependent on RssB, a two-domain response regulator that functions as an adaptor that delivers RpoS to ClpXP for proteolysis. Here we report crystal structures of the receiver domain of RssB both in its unphosphorylated form and bound to the phosphomimic BeF_3_^−^. Surprisingly, we find only modest differences between these two structures, suggesting that truncating RssB may partially activate the receiver domain to a “meta-active” state. Our structural and sequence analysis points to RssB proteins not conforming to either the Y-T coupling scheme for signaling seen in prototypical response regulators, such as CheY, or to the signaling model of the less understood FATGUY proteins.

## 1. INTRODUCTION

Two-component systems mediate signal transduction in bacteria and consist of an environmental cue-sensing histidine kinase that phosphorylates a response regulator composed of either a self-standing “receiver” (REC) domain or a REC domain fused to an effector (EFF) domain^1^. The REC tunes the activity of the EFF domain in response to phosphorylation of a conserved aspartate within the REC. While early studies conceptualized the REC module in terms of a binary logic element (existing in the phosphorylated ON and a non-phosphorylated OFF form), numerous subsequent studies have documented that REC domains exist in a dynamic equilibrium of multiple conformations, with activation sometimes preceding phosphotransfer^2,3^. What has become clear is that phosphorylation-dependent activation often involves small structural changes within self-standing RECs, or oligomerization and closed-to-open transitions within multi-domain REC proteins (reviewed by Gao et al^1^). These changes are often unique to each REC subfamily. Thus, despite a common architectural scaffold, a general mechanistic scheme for REC activation does not hold.

Here we focus on RssB, a two-domain response regulator (**Fig. 1A-B**) and the poorly understood role of its phosphorylation. RssB, a key element of the general stress response in both commensal and pathogenic γ–proteobacteria, is best understood in *Escherichia coli,* where it serves as the only ClpXP adaptor for RpoS, the master regulator of the general stress response^4–7^. RssB delivers RpoS to the ClpXP proteolytic machine for degradation^5,8^. Thus RpoS levels remain low due to rapid turnover when cells are actively dividing. Upon encountering stress, they increase in large part due to inhibition of RssB by so-called anti-adaptors (IraM, IraP, IraD and IraL)^9,10^, favoring competition of RpoS with other promoter specificity subunits for core RNA polymerase^11^. This results in large-scale transcriptional reprogramming^12,13^.

**Figure 1.**
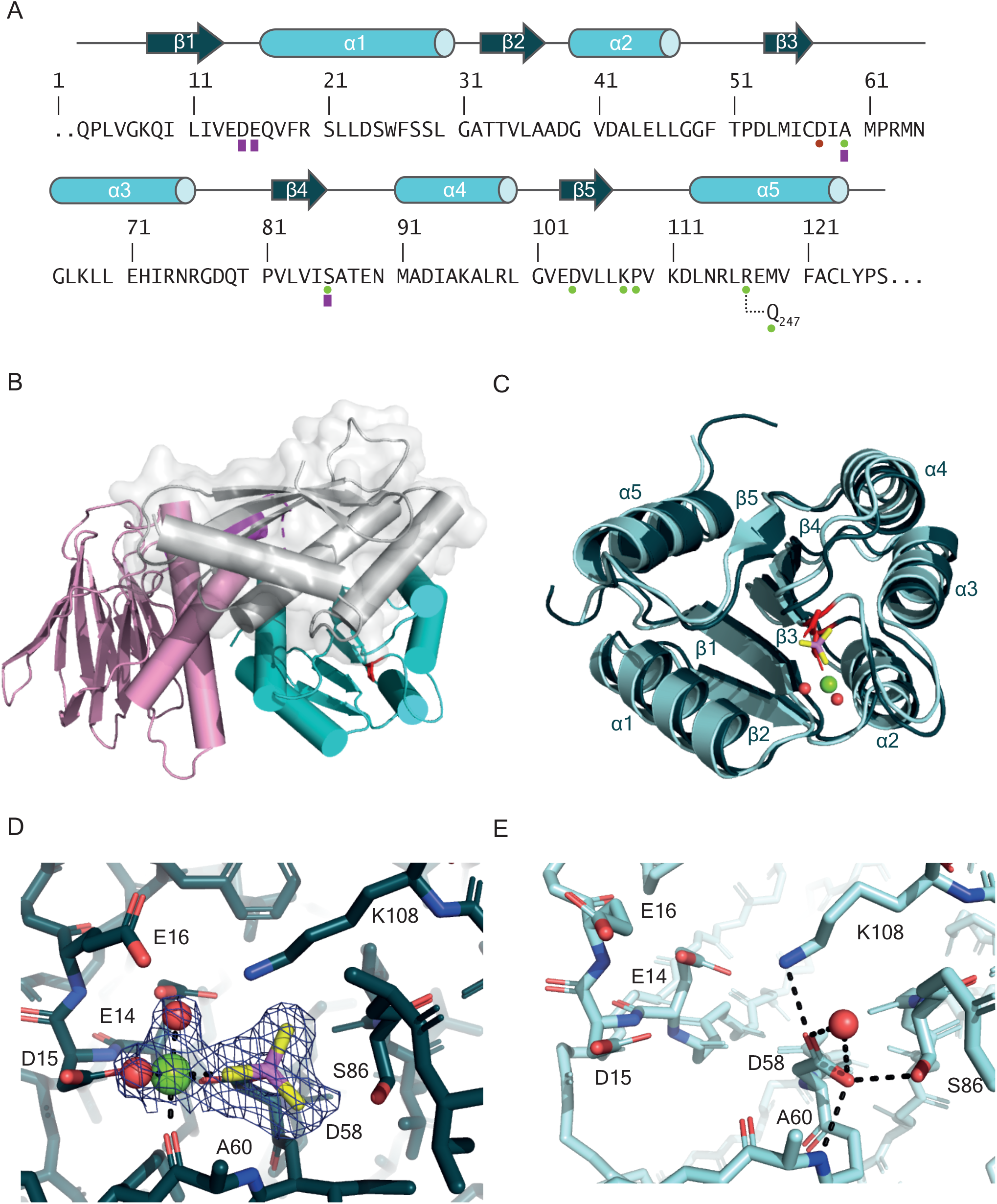
Overall architecture of IraD-bound RssB^D58P^ and RssB^REC^ in the absence/presence of phosphoryl analog, BeF_3_^−^. A. Primary and secondary structure of *E. coli* RssB^REC^. Asp^58^ is indicated with a red circle. Green circles indicate locations at which substitutions result in defects in RpoS degradation^17^. Purple rectangles indicate residues, which together with Asp^58^ constitute the signaling quintet. B. Structure of IraD-bound RssB^D58P^ variant with REC in cyan and the EFF domain in pink. IraD (grey) docks primarily on the 4-5-5 face of RssB^REC^. Residue 58 is shown as red sticks. C. Superposition between apo RssB^REC^ (pale cyan) and RssB^REC^ bound to beryllofluoride (dark teal, rmsd of 0.41 Å). Apo RssB^REC^ also superimposes well (r.m.s.d of 0.28 Å) with a previously reported structure of the same domain in the non-phosphorylated state (PDB ID 6Z4C)^16^. D-E. The phosphorylation site in RssB^REC^ (D) and RssB^REC^•BeF_3_^−^•Mg^2+^ (E). Mg^2+^ is green, while beryllium is magenta and fluoride yellow. A Polder omit map contoured at 3σ (slate mesh) shows clear electron density for the phosphoryl analog, Mg^2+^ and two water molecules (red spheres). In apo RssB^REC^, Asp^58^ hydrogen bonds to Lys^108^, Ser^86^ and a water molecule (red sphere).

RpoS proteolysis is also modulated by phosphorylation of Asp^58^ in RssB^14^, but the few existing reports do not agree on its role^14–18^. This prompted us to solve the crystal structure of the RssB REC domain (RssB^REC^) in its non-phosphorylated form and bound to the phosphomimetic BeF_3_^−^. We find only subtle conformational differences between RssB^REC^ in the presence/absence of Mg^2+^•BeF_3_^−^, suggesting that deletion of the C-terminal domain may partially activate RssB^REC^ even in the absence of a phosphomimic. Such conformational states in which only a subset of residues is at their “active” hydrogen bonding positions have been observed in the context of other response regulators, and have been dubbed “meta-active” states^19–21^. Our analysis also indicates that signaling in RssB is atypical. It lacks canonical Tyr/Thr switch residues, and likely does not conform to the Y-T coupling model seen in classic receiver domains such as CheY^22^. Alternative signaling mechanisms, such as seen in the lesser understood FAT GUY subfamily^23^ are also unlikely to apply, making RssB a response regulator with an unusual mechanism of action.

## 2. RESULTS

### The RssB Phosphorylation Site

We determined the crystal structure of RssB^REC^ (residues 1-129) with an intact phosphoacceptor site (Asp^58^) in its non-phosphorylated state and bound to the BeF_3_^−^ phosphomimetic (**Table 1** and **Figure 1**). This has been found to bind and activate numerous response regulators^24^ and due to the great lability in solution of the natural phosphate-acid linkage, has been widely used in crystallization.

**Table 1.** Crystallographic Data and Refinement Statistics.

|  | **RssB** | **RssB•BeF<sub>3</sub>•Mg<sup>2+</sup>** |
| --- | --- | --- |
| **Data Collection** |  |  |
| X-ray source | Advanced Photon Source<br>24ID-E | Advanced Photon Source<br>24ID-C |
| Space group | P3<sub>1</sub>21 | P3<sub>1</sub>21 |
| Cell dimensions |  |  |
| *a,b,c* (Å)<sup>a</sup> | 46.46, 46.46, 95.85 | 46.33, 46.33, 95.89 |
| $\alpha,\beta,\gamma$ (°) | 90.0, 90.0, 90.0 | 90.0, 90.0, 120.0 |
| Mosaicity (°) | 0.17 | 0.14 |
| Wavelength (Å) | 0.97911 | 0.97918 |
| Resolution (Å) | 95.85-1.73 (1.94-1.85)<sup>a</sup> | 95.89-1.91(1.95-1.91)<sup>a</sup> |
| No. reflections/Unique reflections | 112012/10654 | 90612/9772 |
| Completeness (%) | 99.3/(97.3)\ | 99.5 (93.2) |
| Redundancy | 10.5 (6.4) | 9.3 (6.7) |
| $\langle I/\sigma(I) \rangle$ | 22.3 (2.2) | 21.0 (1.7) |
| R<sub>meas</sub> (all I+ and I-) | 0.058 (0.767) | 0.087(0.992) |
| R<sub>meas</sub> (within I+ / I-) | 0.059(0.784) | 0.088 (0.994) |
| R<sub>pim</sub> (all I+ and I-) | 0.017 (0.291) | 0.028 (0.366) |
| R<sub>pim</sub> (within I+ and I-) | 0.024(0.403) | 0.039 (0.492) |
| CC<sub>1/2</sub> | 0.999 (0.795) | 0.999 (0.690) |
| **Refinement** |  |  |
| Resolution (Å) | 40 – 1.85 | 40.12-1.91 |
| No. reflections, working set | 10624 (977) | 9736 (588) |
| No. of reflections, test set | 1050 (96) | 968 (90) |
| R<sub>work</sub>/R<sub>free</sub> (%)<sup>c</sup> | 19.78/23.03 | 18.2(22.49) |
| No. non-hydrogen atoms |  |  |
| Protein | 868 | 990 |
| Water | 41 | 38 |
| B-factors (Å<sup>2</sup>) |  |  |
| Protein | 48.3 | 41.69 |
| Water | 49.7 | 44.5 |
| Ion/ligand | - | 35.6 |
| R.m.s deviation |  |  |
| Bond lengths (Å) | 0.013 | 0.010 |
| Bond angles (°) | 1.47 | 1.34 |
| Coordinate error (Å)<sup>b</sup> | 0.14 | 0.17 |
| **Ramachandran Plot Analysis** |  |  |
| Preferred (%) | 98.37 | 99.21 |
| Allowed (%) | 1.63 | 0.69 |
| Disallowed (%) | 0 | 0 |
| Rotamer outliers (%) | 0 | 0 |
| PDB ID | 7L9C | 7LCM |

<sup>a</sup> Values in parenthesis are for highest resolution shell
<sup>b</sup> Maximum likelihood estimate

RssB^REC^ has the canonical (βα)_5_ REC fold (**Figure 1C**) and possesses a typical signaling quintet. This includes residues Asp^15^, Glu^16^, involved in Mg^2+^ binding, but also Lys^108^, which reaches towards Asp^58^, and Ser^86^ that stabilizes the Lys^108^ amide both in the absence and presence of beryllofluoride (**Figure 1D-E**). Substitution of Lys^108^ results in negative effects on RssB function *in vivo*^17^ and *in vitro*^16^, consistent with our structures. Similar defects were observed for a Pro^109^ variant (**Figure 1A**). Invariant Pro^109^ mediates a cis-peptide bond in the β5–α5 loop, a conserved feature of many response regulators^25^, which imposes more stringent constraints on stereochemistry and may explain why the *rssB*^P109S^ allele is defective and temperature-sensitive^17^.

The octahedral coordination of Mg^2+^ involves Asp^58^, Asp^15^, the BeF_3_^−^ moiety, two water molecules and the D+2 residue, Ala^60^ (**Figure 1D-E**). Located two residues downstream of Asp^58^, this augments the active site and likely plays key roles in determining the kinetics of REC autodephosphorylation. These can differ by orders of magnitude between different REC domains, and depend heavily on the identity of the D+2 residue^26^. An Ala at position D+2 is atypical for a REC^26^. The K+1 residue, Pro^109^, is invariant, while K+2 (residue 110) is conserved as a branched amino acid (**Supplemental Figure 1)**. These two residues also affect the kinetics of REC phosphorylation/dephosphorylation, but unlike residue D+2, they likely do so via an indirect mechanism that affects REC conformational equilibria^27^. Consistent with this, Val^110^ is located about 12Å away from the phosphorylation site and does not engage in any hydrogen bonds or van der Waals contacts that could link it directly to the active site.

### Signaling by the RssB^REC^ Domain Does not Involve Y-T Coupling

A highly conserved RssB residue is Asp^104^, seen in β5 on the 4-5-5 face (**Figure 2** and **Supplemental Figure 1**). In many response regulators, including CheY, OmpR, NtrC, LytR, NarL, this residue is typically replaced by a tyrosine (e.g Tyr^106^ in CheY)^1^. This tyrosine together with a threonine (Thr^87^ in CheY) is essential for signaling via Y-T coupling. Upon phosphorylation, both Tyr^106^ and Thr^87^ of CheY undergo rotameric changes that propagate from the phosphoacceptor site to the 4-5-5 face^22,28,29^ (**Figure 2C-D**), where many regulators bind, and which, in some systems, supports activation by oligomerization^30^. Thus, the tyrosine sidechain points away from the phosphoryl group in the apo state, and upon phosphorylation, it swing inwards to bury itself in a pocket vacated by the conserved threonine (or serine) in β4. In some cases, it has been found that the threonine equivalent to CheY Thr^87^ gates and engages the entire β4-α4 loop and that the phosphorylation-dependent active conformation of this loop favors burial of Tyr^106^, an extended mechanistic model that has been dubbed T-loop-Y coupling^31^. The presence of an Asp in RssB instead of an aromatic is inconsistent with Y-T coupling. Upon superposition, we also observe no major changes in RssB^REC^ and β5 upon BeF_3_^−^•Mg^2+^ binding (r.m.s.d. of 0.41 Å). The 4-5-5 face, the locus of the largest differences between ON and OFF conformations in REC domains, is remarkably similar, perhaps due to the partial activation of RssB^REC^ in this truncated form. However, significant conformational differences are observed between the 4-5-5 face of RssB^REC^ and of full-length RssB^D8P^ due to tilting of α5 as well as motion of the loop preceding it (**Figure 2A**). This points to this face of RssB^REC^ being plastic and subject to regulation.

**Figure 2.**
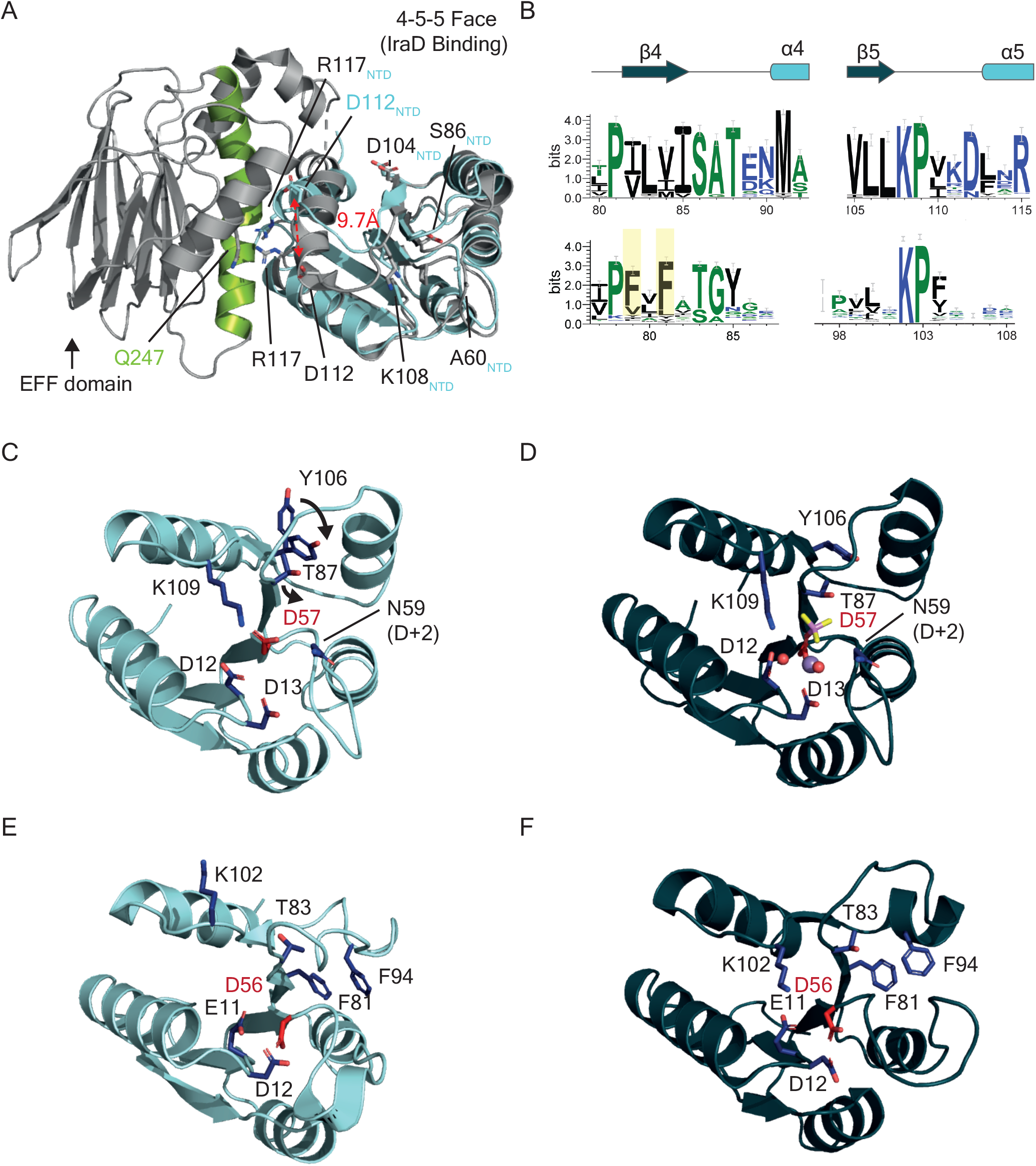
RssB does not employ Y-T coupling nor SdrG-like signaling. A. Superposition of RssB^REC^ (pale cyan) and full-length RssB^D58P^ mutant (PDB ID 6OD1^40^, grey). The signaling helix, site of class II mutations as defined by Gottesman and coworkers is colored green. Asp^104^ is shown as sticks. B. Sequence conservation around the PvL[VIM]ISAT motif in the RssB family and equivalent region in SdrG. Residues are colored by charge/hydrophobicity. Substitution of boxed SdrG residues lead to loss of function^23^. C-D. Structures of *E. coli* CheY in OFF state (PDB ID 3CHY)^25^ and ON state (PDB ID 1FQW)^44^. Phosphoacceptor is shown in red. The ON state structure was solved by NMR and contains a Mn^2+^ rather than a Mg^2+^ ion. Two Tyr^106^ rotamers were observed. Overall rmsd after superposition of the ON/OFF states is 0.54 Å. E-F. *Sphingomonas melonis* SdrG in the OFF state (PDB ID 5IEB)^23^ and ON state (PDB ID 5IEJ)^23^. Phosphorylation site is shown in red. Structures were solved using NMR, and the positions of the phosphomimetic and Mg^2+^ were not determined experimentally. For simplicity they were not modelled. Overall superposition r.ms.d. is 1.68 Å.

### Comparison to FAT GUY Response Regulators

The lack of Y-T coupling is unusual, but not without precedent. In CheY2, the role of Tyr^106^ is taken by Phe^59^ located in the β3-α3 loop^32^, while in NtrC, the rotameric state of the tyrosine is uncorrelated with the global conformational transition during activation^33^. In the NarL subfamily, the tyrosine exists but is constitutively pointing inward regardless of phosphorylation state^34,35^, suggesting it may not be involved in allostery. In response regulators of the FATGUY family, such as SdrG, activation occurs instead via stabilization of the phosphoryl-aspartate via hydrogen bonding with Lys^102^ (Lys^108^ in RssB) and Thr^83^ (Ser^86^ in RssB), which occupies the same position within β4 as the conserved CheY threonine involved in Y-T coupling (**Figure 2**). The rotation of Thr^83^ ensues in remodeling of several phenylalanines, in and around a highly conserved PFXFAT^83^G[G/Y] motif making up the β4 strand (**Figure 2B,E, F**)^23^. Activation also leads to structuring of the α4 helix and changes in the loop C-terminal to the phosphoacceptor site. In RssB^REC^ and RssB^REC^• BeF_3_^−^•Mg^2^ and the recent structure of IraD-bound RssB^D58P^ (**Figure 2A**), we observe that the equivalent Lys^108^ points towards the phosphoacceptor location. This is usually associated with an ON state and is contrasted to the OFF conformer, in which Lys^108^ points away (**Figs. 2E** and **F**). However, neither of the remodeled conserved FATGUY residues is present in the consensus RssB sequence. Instead, they are replaced by a highly conserved PvL[VIM]ISAT^88^ motif spanning β4 and the β4-α4 loop (**Figure 2B**), with the phenylalanines being replaced by hydrophobes (Val, Ile, Met). Notably, Phe^79^, Phe^81^ and Phe^94^ lead to functional SdrG defects *in vitro* and *in vivo*^23^. While conservation of the residues in β4, part of the hydrophobic core, may be due to scaffolding reasons, Leu^83^ and Ile^85^ interact with Cys^57^ next to the phosphorylation site, and may relay phosphorylation status to more distant sites, such as the β4-α4 loop. This loop contains conserved residue Thr^88^ (also dubbed as T+2 according to the nomenclature of Bourret and colleagues), which modulates dephosphorylation rates in family-specific ways^36^. In both of our structures, Thr^88^ hydrogen bonds to D+1 residue Ser^86^ and Asn^90^. As a consequence, there are minimal differences between the β4-α4 loops in apo REC and the REC~P-like form. We cannot rule out that this may be due to using truncated protein constructs for crystallization.

### The Importance of Helix α5

Temperature factor analysis confirms that the β3-α3 loop C-terminal to Asp^58^ displays increased dynamics in RssB^REC^ in the absence of phosphorylation (**Supplemental Figure 2A**). Surprisingly, helix α5 part of the central helical bundle to which both the N-terminal and C-terminal domains contribute in full-length RssB (**Figures 2A**) undergoes only subtle changes, notably the rotameric conformations of Lys^111^, Arg^117^, and Phe^121^ in α5 (**Supplemental Figure 2C-F**).

The interactions of α5 residue Arg^117^ attracted our attention for four reasons. First, in RssB^REC^•BeF_3_^−^•Mg^2+^, the amide of Arg^117^ hydrogen bonds to the carbonyl of Arg^115^, while in RssB^REC^, Arg^117^ is stabilized by contacts with the π systems of Phe^121^ and Trp^26^ (**Supplemental Figure 2C, D**). Secondly, the tilting of α5 we observe in the RssB^REC^ structure relative to IraD-bound RssB^D58P^ would cause a clash with α9 in the closed RssB structure (**Figure 2A**). α9 is, incidentally, the site of mutations (class II mutations as defined by Gottesman and coworkers and highlighted in green **Supplemental Figure 2B**) that activate RssB in the absence of phosphorylation, bypassing the need for acetyl phosphate^17^. Thirdly, in IraD-bound RssB^D58P^, Arg^117^ interacts with Gln^247^, and this interaction dominates the REC-EFF interface (**Supplemental Figure 2B**). Last, but not least, an alanine substitution of Arg^117^ results in loss of interaction with RpoS in the absence of acetyl phosphate, although degradation (and, presumably substrate binding) is only partially compromised *in vitro* in the presence of acetyl phosphate^16^. This points to Arg^117^ serving as a switch that modulates substrate binding.

## DISCUSSION

As of November 2020, PFAM^37^ lists over 324,000 REC sequences, and the Protein Data Bank contains many structures of REC domains, mostly in the unphosphorylated state. With such a bounty of structural information, one would expect a thorough understanding of how signaling takes place in REC proteins. This is not so, since REC domains are often stringed to a variety of domains supporting different REC-EFF contacts, structures of REC~P-like states are relatively rare, and reaction kinetics are strongly influenced by sub-family specific residues (e.g D+2, T+2, but not only) outside the conserved architecture of the phosphorylation site^26^. Our structure of apo RssB^REC^ reveals a meta-active conformation that resembles to a great extent RssB^REC^•BeF_3_^−^•Mg^2+^. The structural differences between RssB^REC^ within IraD-inhibibted RssB^D58P^ and RssB^REC^•BeF_3_^−^•Mg^2+^, are modest but involve the β3–α3 (**Supplemental Figure 3**) in agreement with the general structural framework of REC activation^38,39^. While a full understanding of RssB mechanism will require the structure determination of full-length RssB and RssB~P bound to its various partners, our work allows us to reconsider the role of phosphorylation in the RpoS general response. It has generally been viewed that phosphorylation by either kinases or small phosphodonors promotes RpoS binding and turnover by ClpXP^15,17^. More recent studies have proposed that, on the contrary, phosphorylation reduces RssB affinity for RpoS, and consequently the rate of RpoS degradation. This model was based on equilibrium dissociation constant determination for core RssB-RpoS complexes comprised of RssB^REC^ and region 3 of RpoS (residues 163-220), and a less than two-fold decrease in the affinity in the presence of acetyl phosphate^18^. This model is at odds with multiple observations, including the requirement for acetyl phosphate in promoting RpoS-RssB^17^ and RpoS-RssB-ClpXP complex formation^5^, and stimulation of RpoS degradation in an *in vitro* system^17,40^. It is however easily explained by our determination that removing the EFF domain partially activates RssB^REC^ independent of phosphorylation, obfuscating the effects of phosphoryl transfer on the full-length proteins.

We suggest that RssB does not conform to the YT-coupling model for signaling seen in many response regulators, nor to the signaling mechanism used by FATGUY proteins during the alphabacterial stress response. RssB deregulation achieved by a single amino acid substitutions of Arg^117^ in helix α5 severely impairs function^16^. This, together with the major conformational differences between IraD-bound RssB^D58P^ (in an OFF state) and RssB^REC^ point to helix α5 as key for signal propagation to the EFF domain. Consistent with this, mutations in the interdomain linker C-terminal to α5 (W143R, P150S, E135A E136A E137A) have also been demonstrated to affect the adaptor function of RssB or its ability to be regulated by anti-adaptors^17,40^. Without doubt, further understanding of the regulatory RssB system will require a multi-pronged approach, including determination of structures of full-length RssB and RssB complexes, which, more than two decades since the discovery of RssB’s role as a RpoS regulator^5,6,41^, remain eagerly awaited.

## METHODS

### RssB^NTD^ Production and Crystallization

RssB^NTD^ was purified as a hexahistidine tag 6His-RssB(1-129) fusion a succession of nickel-affinity chromatography, tag cleavage and dialysis followed by gel filtration on a Superdex 75 Hi Load column 16/60 (GE Healthcare) pre-equilibrated with 20mM Tris-HCl pH 8, 200mM NaCl, 10mM MgCl_2_, and 2mM TCEP. The sample was crystallized against 0.2M lithium sulfate, 22% PEG 3,350, and 0.1M Tris-HCl pH 8.8 in sitting drop format. Cryoprotection was achieved by supplementation of the well solution with 25% ethylene glycol. RssB^NTD^-BeF_3_^−•^Mg^2+^ was crystallized in conditions similar to those identified for RssB^NTD^.

### Structure Determination and Refinement

Diffraction data was processed using XDS^42^ and phased using molecular replacement with the homologous domain in PDB ID 6OD1 in Phaser^43^. Iterative refinement and building were performed in PHENIX and Coot. A final refinement cycle was carried out using Refmac with PDB-REDO. Diffraction images have been deposited in the SBGrid Data Bank as dataset number 815 (doi:10.15785/SBGRID/815) and 818 (doi:10.15785/SBGRID/818). PDB IDs are listed in Table 1.

## ACKNOWLEDGEMENTS

We thank Drs. S. Gottesman (NIH) for critical reading of the manuscript. Diffraction data were collected at sector 24ID at the Advanced Photon Source. This research was supported by grant R01GM121975 (National Institutes of Health) and a Salomon Research Award (Brown University) to A.M.D.

## DATA AVAILABILITY STATEMENT

Data and reagents are available from the corresponding author upon reasonable request.

## AUTHOR CONTRIBUTION

J.S. purified protein and obtained crystals, and J. Son collected diffraction data and refined structures. A.M.D designed the overall study, collected diffraction data, solved and refined structures, and wrote the manuscript with figure contributions from C.B.

