## Supplementary Information for "Phospho-dependent Signaling during the General Stress Response by the Atypical Response Regulator and ClpXP Adaptor RssB"

**FOR**

**Supplementary Figure 1. Consensus RssB sequence based on an alignment of 97 RssB sequences.** Residues are colored by charge.

**Supplementary Figure 2. Context-dependent interactions of Arg$^{117}$.**

A. Plot of the isotropic temperature (B) factor for $RssB^{REC}$ and $RssB^{REC}$•$BeF_3^-$•$Mg^{2+}$.

B. View of the central helical bundle in IraD-bound $RssB^{D58P}$. Colored in green are residues shown previously to bypass the need for phosphorylation. These pack against residues in α5 such as Tyr$^{125}$, which in turn interacts with Trp$^{143}$ within the interdomain linker. Mutation of Trp$^{143}$ compromises regulation of RpoS proteolysis by IraD and IraP[17]. Arg$^{117}$ is hydrogen bonded to Gln$^{247}$ (inset).

C-D. View of the residues highlighted in C in the context of $RssB^{REC.}$ View of α1 is obscured by α5, and a better view of Arg$^{117}$ and Trp$^{26}$ is shown in D.

E-F. View of the residues highlighted in C in the context of $RssB^{REC}$•$BeF_3^-$•$Mg^{2+}$. The $BeF_3^-$ moiety is shown in yellow/magenta and $Mg^{2+}$ in green. Water molecules have been omitted for simplicity. View of α1 is obscured behind α5, and a better view of non-interacting Arg$^{117}$ and Trp$^{26}$ is shown in F.

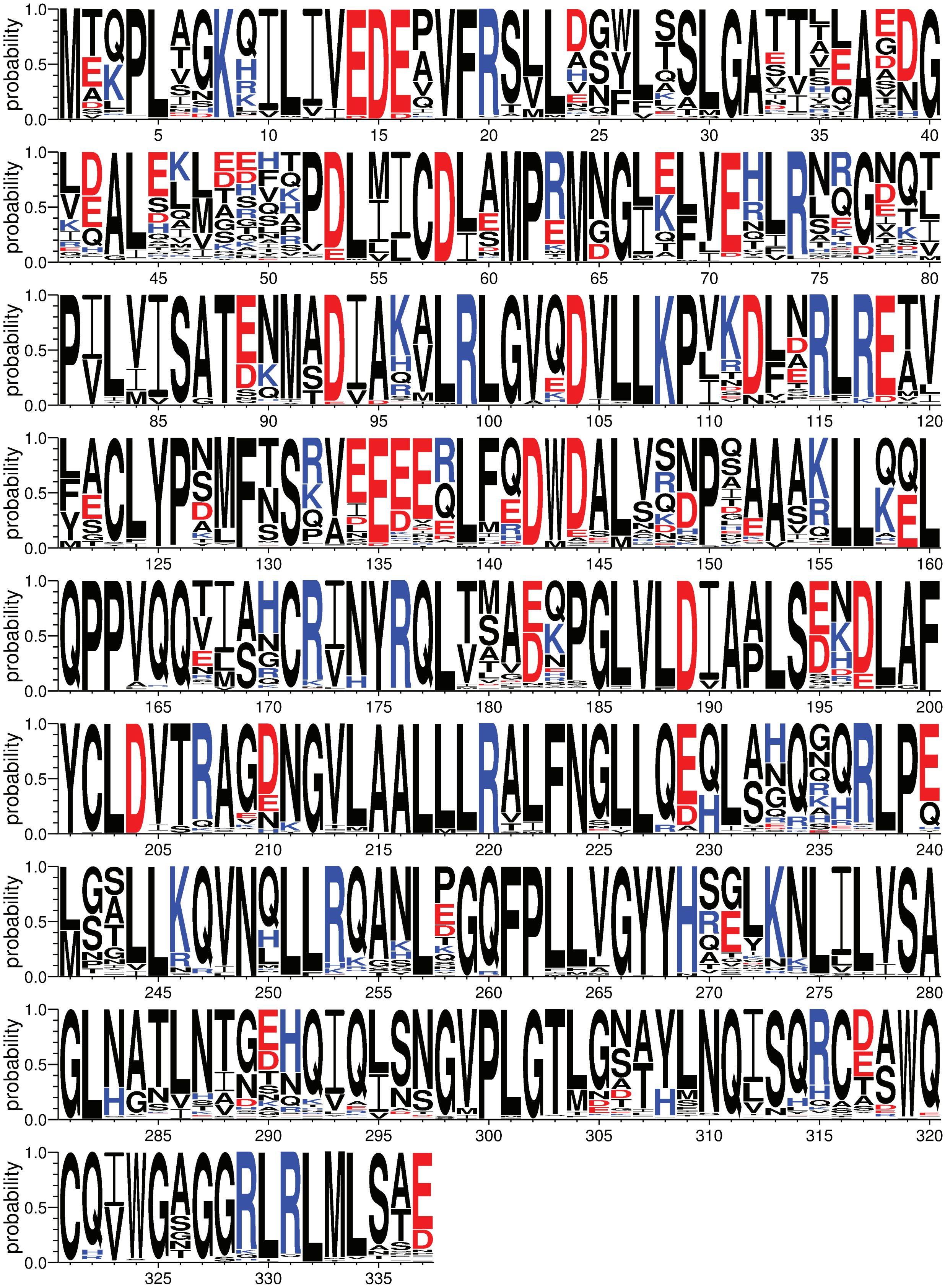

Supplemental Fig. 1

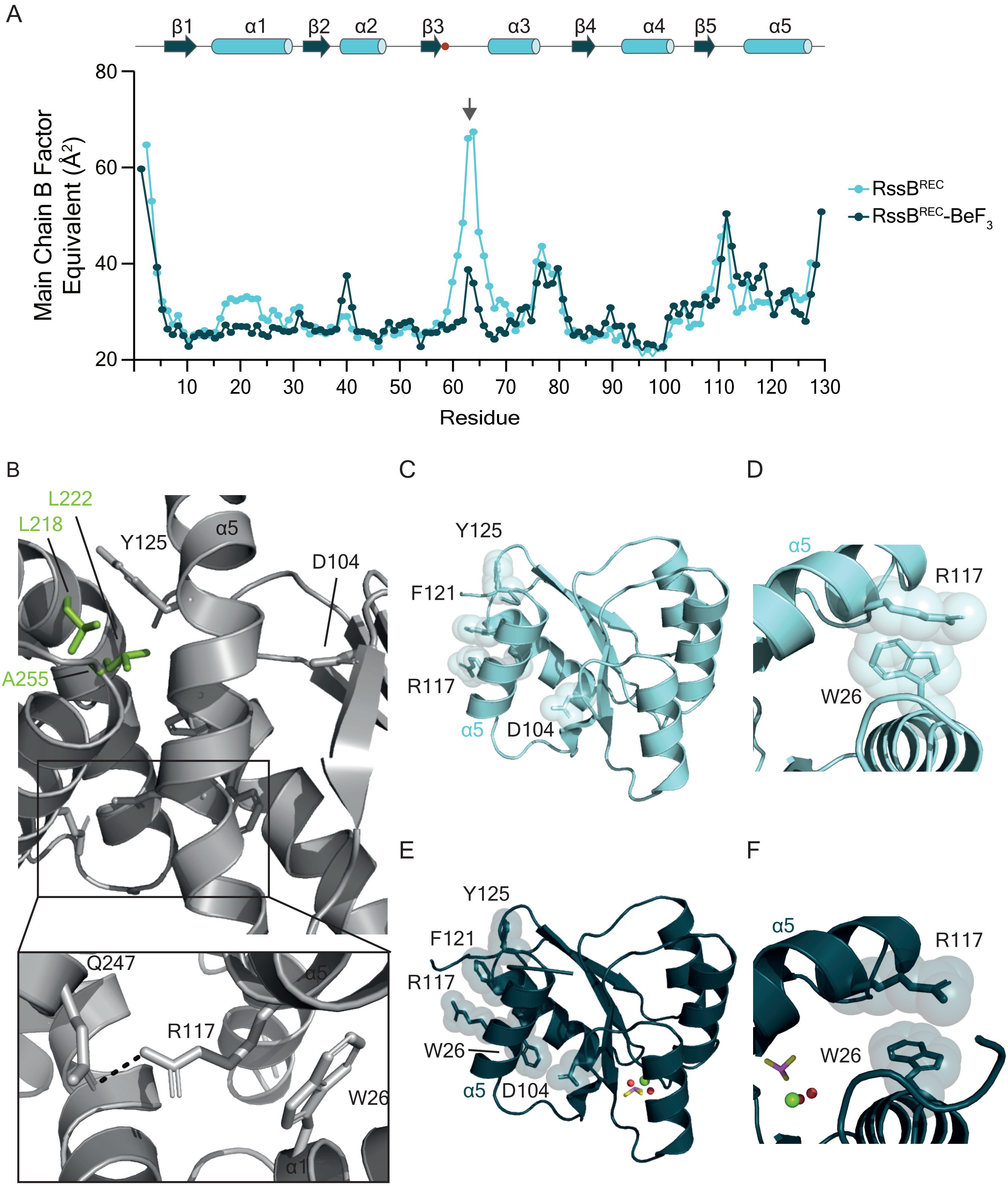


Supplemental Figure 2
